# TuMV triggers stomatal closure but reduces drought tolerance in Arabidopsis

**DOI:** 10.1101/2020.08.03.235234

**Authors:** Carlos Augusto Manacorda, Gustavo Gudesblat, Moira Sutka, Sergio Alemano, Franco Peluso, Patricio Oricchio, Irene Baroli, Sebastián Asurmendi

## Abstract

i)

In this work the effects of TuMV infection on stomatal closure and water balance were studied in Arabidopsis. Thermal imaging analyses showed that TuMV-infected plants had consistently higher foliar temperature than mock treated controls. Non-destructive time-course experiments revealed that this differential phenotype was stable during both daytime and nighttime. This effect was due to reduced gas exchange in TuMV-infected plants, as observed through stomatal conductance and stomatal aperture assays in systemic leaves. Measurements of daily water consumption and initial dehydration rate further proved that TuMV infection reduced water loss. Salicylic acid (SA) and abscisic acid (ABA) contents were increased in TuMV-infected plants. In addition, the expression of ABI2, involved in ABA signaling, was enhanced, and ABCG40 (required for ABA transport into guard cells) was highly induced upon TuMV infection. Hypermorfic *abi2-1* mutant plants, but no other ABA or SA biosynthetic, signaling or degradation mutants tested abolished both stomatal closure and low stomatal conductance phenotypes caused by TuMV. Therefore, not overall ABA levels, but localized differences in ABA import and perception in guard cells, are likely to be responsible for stomatal closure observed under TuMV infection. Plants simultaneously subjected to drought and viral stresses showed higher mortality rates than their mock-inoculated drought stressed counterparts, consistent with down-regulation of drought-responsive gene RD29A, both in short and long day conditions. Our findings indicate that in spite of stomatal closure triggered by TuMV, additional phenomena cause compromised drought tolerance of TuMV-infected Arabidopsis plants.

**Significance statement:** Characterization of the physiological responses controlling plant water management under combined stresses and the genes behind them is important in the current climate change scenario, which poses multifaceted challenges to crops. We found that TuMV infection induced ABA and SA accumulation and stomatal closure in Arabidopsis, alongside with overexpression of ABCG40 (the transporter of ABA to guard cells), whereas the dehydration-responsive gene RD29A was downregulated, concomitantly with increased plant susceptibility to drought stress under infection.

## iii) Introduction

### Viral infections and drought tolerance

Phytoviruses are a common cause of agricultural losses worldwide (Loebenstein, 2010). Compatible viral infections often diminish crop value, affecting important traits (Matthews *et al.*, 2002). Therefore, it is widely accepted that viral infections are harmful to the host. However, this paradigm has been challenged in recent years, as more evidence piled up revealing that the outcome of viral infections depends strongly on environmental conditions (Ramegowda and Senthil-Kumar, 2015), an increasingly important issue in view of ongoing global climate change (Bergès *et al.*, 2018; Hily *et al.*, 2016; Grimmer *et al.*, 2012). Under certain environmental circumstances like drought, plant viruses (as reported for some animal viruses (Ryan, 2004)), can be in fact conditional mutualistic partners of plants (Xu *et al.*, 2008; Hily *et al.*, 2016). Therefore, whereas sometimes detrimental to the plant’s fitness, viruses could under certain environmental conditions, allow the plant to better tolerate certain abiotic stresses (Carr *et al.*, 2018). These tolerant phenotypes are associated with transpiration rates that are either lower (Xu *et al.*, 2008; Aguilar *et al.*, 2017) or higher (Aguilar *et al.*, 2017; Westwood *et al.*, 2013), due to the fact that the impact of viruses on host stomatal development and function depends on each pathosystem. In compatible interactions, viruses have been shown to alter stomatal development (Hall and Loomis, 1972; Murray *et al.*, 2016; Aguilar *et al.*, 2017) or reduce stomatal aperture (Lindsey and Gudauskas, 1975; Grimmer *et al.*, 2012). However, the molecular mechanisms by which pathogens manipulate stomatal development or aperture are not well understood (Melotto *et al.*, 2017).

### Viral infections and plant hormone responses

Paramount in the regulation of stomatal opening is the phytohormone abscisic acid (ABA), involved in the response and tolerance to diverse abiotic stresses, including drought (Vishwakarma *et al.*, 2017; Ma *et al.*, 2018). Salicylic acid (SA) also plays an important role in various plant developmental processes and responses to abiotic and biotic stresses (Raskin, 1992; Vlot *et al.*, 2009; Maruri-López *et al.*, 2019) including stomatal defense (Melotto *et al.*, 2017; Wang *et al.*, 2018; Klessig *et al.*, 2018). ABA has been more often studied in its role in abiotic stress response whereas SA is viewed mainly as acting in biotic (including anti-viral) interactions; however, a growing corpus of evidence points out to a cross-talk between these hormones under both types of stresses (Wang *et al.*, 2018; Alazem and Lin, 2015; Alazem, Kim, *et al.*, 2019; Pasin *et al.*, 2020). Jasmonic acid (JA) is another major plant defense hormone (Pieterse *et al.*, 2012; Alazem and Lin, 2015) which, together with SA, orchestrates the defense response to a diverse range of pathogens. Both SA and JA act mainly in an antagonistic way, with SA being a stronger repressor of JA pathway than vice-versa (Vlot *et al.*, 2009; Hickman *et al.*, 2019). JA derivatives were also reported to play a role in stomatal closure (Melotto *et al.*, 2017). The interaction between SA and JA can in turn be modulated by ABA and the cross-talk between ABA and SA is also bidirectional (Vlot *et al.*, 2009). In recent years, the interconnected nature of hormone responses to pathogens has been increasingly emphasized (Alazem and Lin, 2015; Vlot *et al.*, 2009; Klessig *et al.*, 2018; Collum and Culver, 2016).

The hormone profile following viral infections is not homogeneous and depends on the specific pathosystem under study (Collum and Culver, 2016). Several plant-virus interactions display a strong ABA response (Whenham *et al.*, 1986; Alazem and Lin, 2017; Alazem and Lin, 2015) whereas in occasions compatible viral infections can disrupt hormonal balance, resulting in simultaneous induction of ABA and SA pathways, which were traditionally viewed as antagonistic, as in the case of BaMV and CMV infections (Alazem *et al.*, 2014).

Among plant viruses Potyvirus is the largest genus, causing significant losses in a wide range of crops (Revers and García, 2015). Due to their importance, potyviruses are between the most studied phytoviruses and their study covers many aspects of plant virology. However, only a few studies were conducted to date to analyze the impact of potyviral infection under abiotic stress (Revers and García, 2015; Prasch and Sonnewald, 2013). TuMV is one of the best studied viruses of its genus, partially due to its ability of infect the model plant Arabidopsis as well as dozens of species of economic importance (Walsh and Jenner, 2002). TuMV has been shown to interfere with signaling of hormones such as ethylene, ABA and SA (Casteel *et al.*, 2015; Poque *et al.*, 2018; Sánchez *et al.*, 2015) and triggers early accumulation of SA-responsive senescence associated genes (Manacorda *et al.*, 2013).

Here we studied the effect of TuMV infection on water management of Arabidopsis at the level of the stomatal response. TuMV induced partial closure of Arabidopsis stomata as well as increases in SA and ABA content. Genes responsible for ABA biosynthesis and catabolism were downregulated, but genes involved in ABA signaling, and particularly in ABA transport to guard cells, were highly induced by TuMV. ABA, SA and JA loss of function biosynthetic mutants did not abolish the phenotype, whereas *abi2-1* hypermorphic ABA signaling mutants exacerbated it to a mild degree. These results indicated that not overall ABA levels, but localized differences in ABA import and perception in guard cells, could influence the closure of stomata under infection. In spite of the shift in hormonal balance, TuMV increased the susceptibility of Arabidopsis plants to drought, in line with the down-regulated transcription of the classic reporter of drought tolerance gene RD29A, both in short day (SD) and long day (LD) conditions. The increased susceptibility to drought of *A. thaliana* plants infected with TuMV is consistent with previously reported findings (Prasch and Sonnewald, 2013) and its occurrence under two different growth conditions in our study points out to a rather robust phenomenon.

## Results

### TuMV induces stomatal closure and lowers water loss in Arabidopsis

To investigate whether TuMV alters rosette transpiration in Arabidopsis, we inspected TuMV-infected plants non-destructively by infrared (IR) imaging (Costa *et al.*, 2013; Pesti *et al.*, 2019). Figure S1a shows the typical visible phenotypic outcome of TuMV infection (UK1 strain) under SD conditions (Manacorda and Asurmendi, 2018). TuMV-infected plants showed a T_leaf_ increase between 0.5-1°C relative to the mock-inoculated controls after 2 weeks post-inoculation (DPI) (Figure S1b). It is known that JPN1 and UK1 strains of TuMV cause differential phenotypical and molecular responses in Arabidopsis (Sánchez *et al.*, 2015; Manacorda *et al.*, 2013). Additionally, it has been observed that, as viral infections progress over time, changes occur in several important traits such as viral accumulation, viral-induced accumulation of metabolites and defense-related transcripts, and plant morphology, at a rate that is specific for each pathosystem (Alazem *et al.*, 2017; Manacorda *et al.*, 2013; Wang *et al.*, 2011; Bazzini *et al.*, 2011; Manacorda and Asurmendi, 2018). Therefore, the use of the time-course methodology is useful for obtaining a more accurate insight (Alazem *et al.*, 2017; Alazem and Lin, 2015). Taking advantage of the non-destructive nature of IR imaging, we performed a time-course observation of TuMV-induced T_leaf_ increase in Arabidopsis for JPN1 and UK1 strains (Figure 1). After 8 DPI both strains caused statistically significant increases in T_leaf_ relative to mock treated plants, while no differences were observed between strains throughout the experiment (Figure 1c). Because T_leaf_ is routinely taken as a proxy of the stomatal conductance (*g_s_*) (Costa *et al.*, 2013; Merlot *et al.*, 2002), and several plant-virus interactions affect both water balance and *g_s_* (Xu *et al.*, 2008; El Aou-ouad *et al.*, 2017), we investigated whether TuMV modifies *g_s_* in Arabidopsis. We observed that *g_s_* values were statistically significantly lower in TuMV-infected samples, without differences between strains (Figure 1d). Therefore, T_leaf_ in Arabidopsis negatively correlated with measured *g_s_*, as expected. Given that both TuMV strains provoked similar IR and *g_s_* phenotypes, we selected the UK1 strain to continue our work.

**Figure 1.**
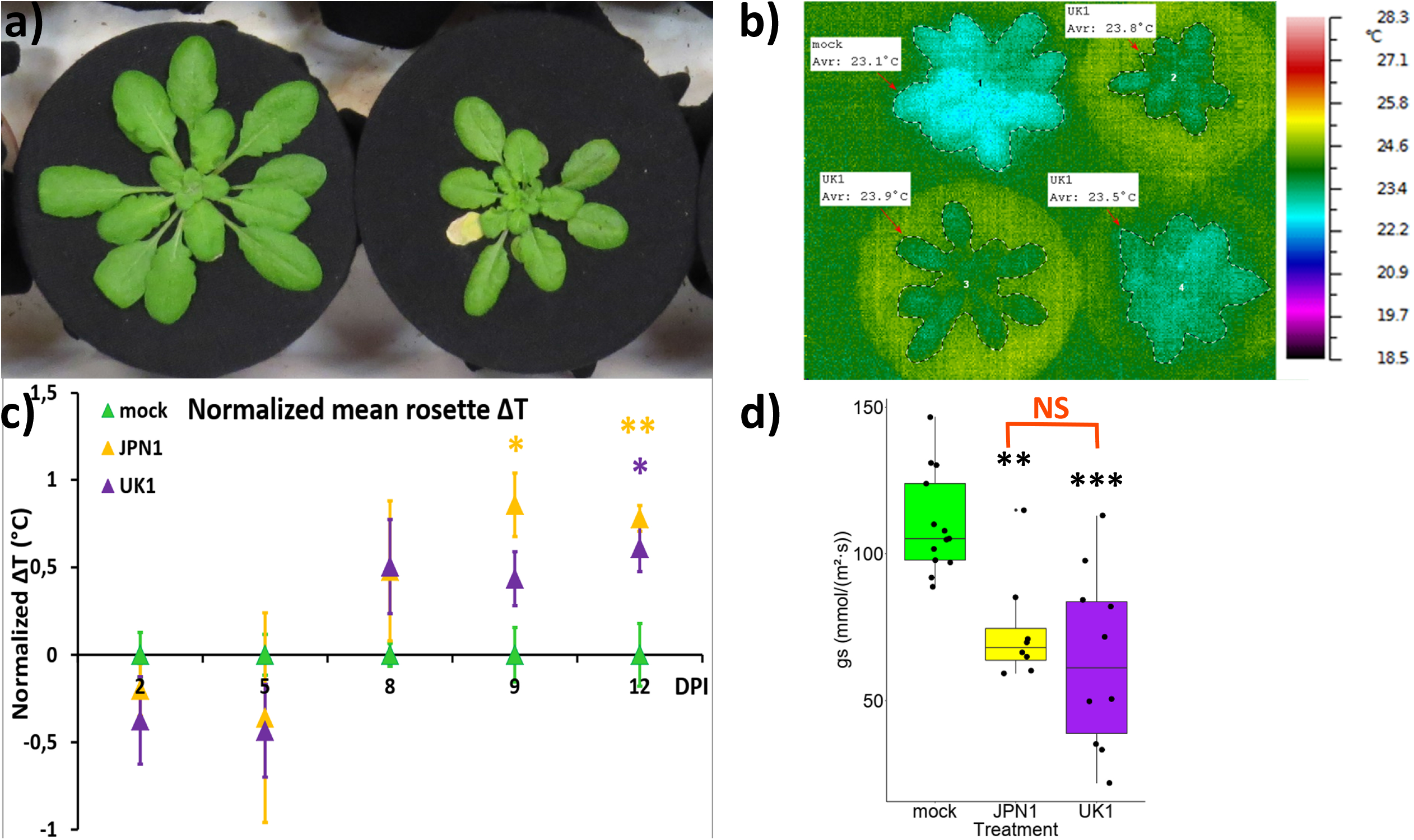
Both JPN1 and UK1 TuMV strains similarly increase T_leaf_ and reduce stomatal conductance in Arabidopsis. ***(a)*** Representative phenotypes of mock-inoculated controls (to the left) and TuMV (UK1)-infected (to the right) in plants growing in soil covered by breathable cloth at 12 DPI. ***(b)*** Representative thermal image (false-color scale to the right) showing T_leaf_ (average rosette temperature) in TuMV (UK1) and mock plants. ***(c)*** ΔT_leaf_ normalized data of TuMV JPN1 and UK1 (5 ≤ n ≤ 9 per group). Asterisks indicate statistically significant differences relative to the controls. ***(d)*** *G*_*s*_ values for both TuMV strains (JPN1 and UK1) and mock plants (box-and-jitter plot). Tukey’s multiple comparisons test. 8 ≤ n ≤ 13 per group.

Microscopy imaging observation confirmed that TuMV induced stomatal closure in systemic leaves (Figure 2). The reduction in stomatal aperture was nearly 40% in average (p < .0001). Whereas stomatal aperture is determined, among other factors, by light intensity, pathogens such as bacteria can trigger stomatal closure under bright daylight (Melotto *et al.*, 2017; Gudesblat *et al.*, 2009), overriding the natural circadian rhythm of stomatal movement. During the night, stomata experience a partial closure as part of this cycle (Caird *et al.*, 2006; Lasceve *et al.*, 1997). We next investigated whether differences in stomatal aperture between treatments were maintained during nighttime (when stomata of mock-inoculated controls are expected to be already closed relatively to daytime). A hydroponically-grown setup optimized for Arabidopsis was employed (Conn *et al.*, 2013). This technique ensures equal access of roots to water and nutrients along the entire experiment, while allowing a more packed scheme for simultaneously imaging several plants without sacrificing pot size (an issue that should be considered when plants are allowed to grow during several weeks). The phenotypes triggered by TuMV could be reproduced in hydroponically-grown plants (Figure S2a). In a time-course experiment between 2-15 DPI, infrared images showed a similar pattern than observed in soil-grown plants (Figure 1c, Figure S2b), both in timing of appearance of significant T_leaf_ differences and in their magnitude. Afterwards, infrared images were automatically taken encompassing periods of late daytime, dusk, the entire nighttime, sunrise and early daytime, simulated in the growth chamber (Figure S2c-e). Drops in T_leaf_ for all plants during nighttime reflect the temperature drop in the growth chamber. T_leaf_ differences between treatments, however, did not change through nighttime, either considering all measurements (Table S1, interaction treatment:time with P = 0.11971), or considering periods of day, (dawn, day, dusk and night, Table S2. Interaction treatment:period with P = 0.73218). Expectedly, in each analysis both time/period and treatment affected T_leaf_ (0.0001<P< 0.01) due to lower programmed chamber temperature during nighttime and TuMV treatment, respectively. Although non-statistically significant, we did detect differences in the average ΔT between treatments between daytime and nighttime (ΔT = 0.45 °C and ΔT = 0.31 °C, respectively, Table S3), indicating that at night, differences between treatments are somewhat lower, perhaps due to stomata of TuMV-infected plants being already partially closed during daytime. The expected endogenous behavior of stomata at night with a low peak at midnight and then a slow increase in predawn hours (Caird *et al.*, 2006) could also be detected for both treatments. Therefore, infection with TuMV reduces stomatal aperture throughout the day, during both daytime and nighttime, and does not seem to interfere with the natural circadian rhythm of stomatal movement. Afterwards, taking advantage of the hydoponic setup, we carried on experiments of whole-plant daily relative water consumption. Firstly, we verified through gs measurements, that TuMV impaired water loss through stomata in hydroponically-grown plants (Figure S3). Then, we analyzed daily relative water consumption, which was reduced by TuMV (Figure 3a). In addition, experiments of initial dehydration rate of detached rosettes showed that evapo-transpiration rates of rosettes (Figure 3b-c) were reduced by TuMV.

**Figure 2.**
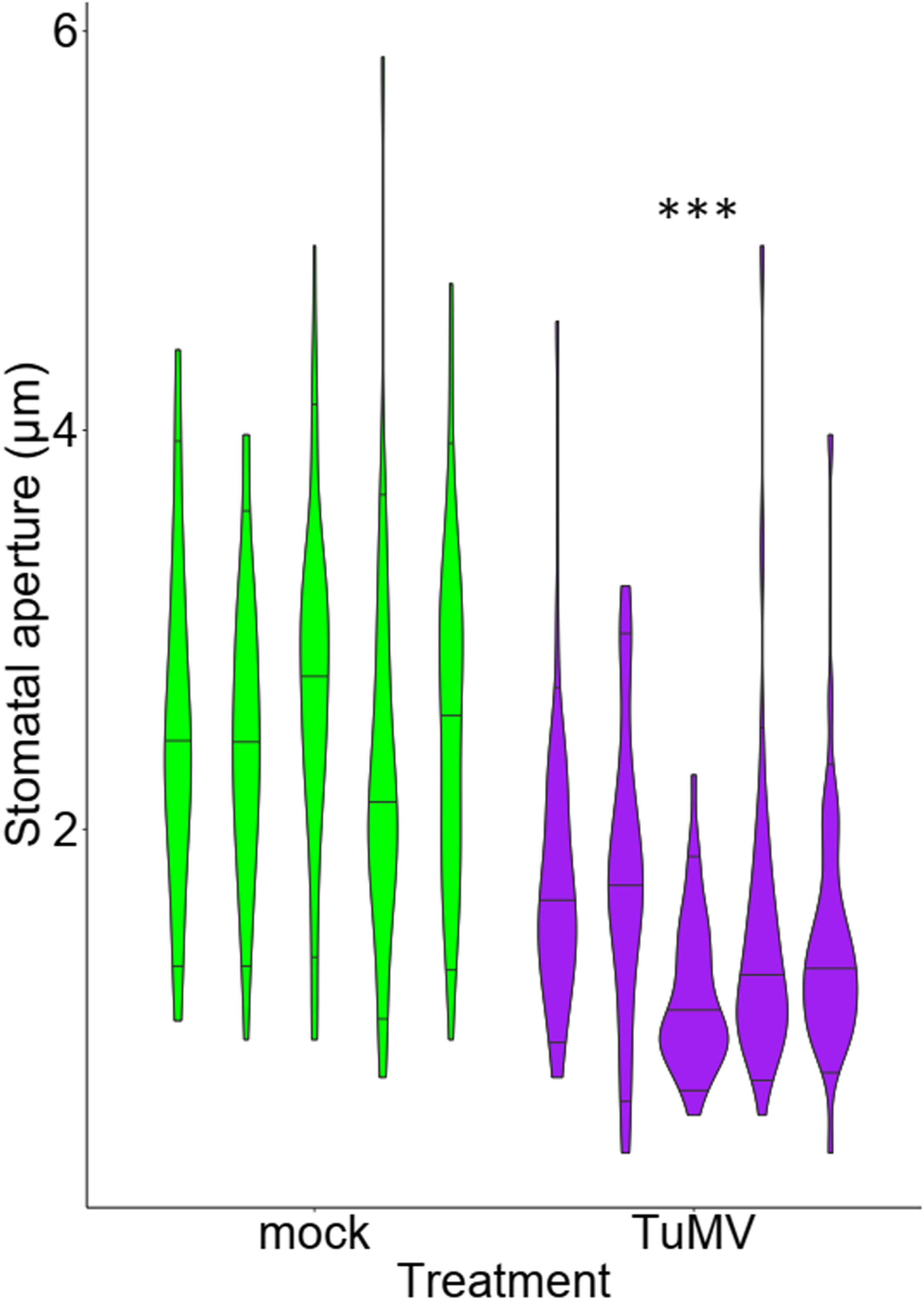
TuMV decreases stomatal aperture in systemic Col-0 leaves. Stomatal aperture in control and TuMV-infected plants at 12 DPI. Violin plots show 0.05, 0.5 and 0.95 quantiles for stomatal aperture data (n = 40) from each individual. Data from two independent experiments showed similar results. Results of one representative experiment are shown, n = 5 for each treatment group.

**Figure 3.**
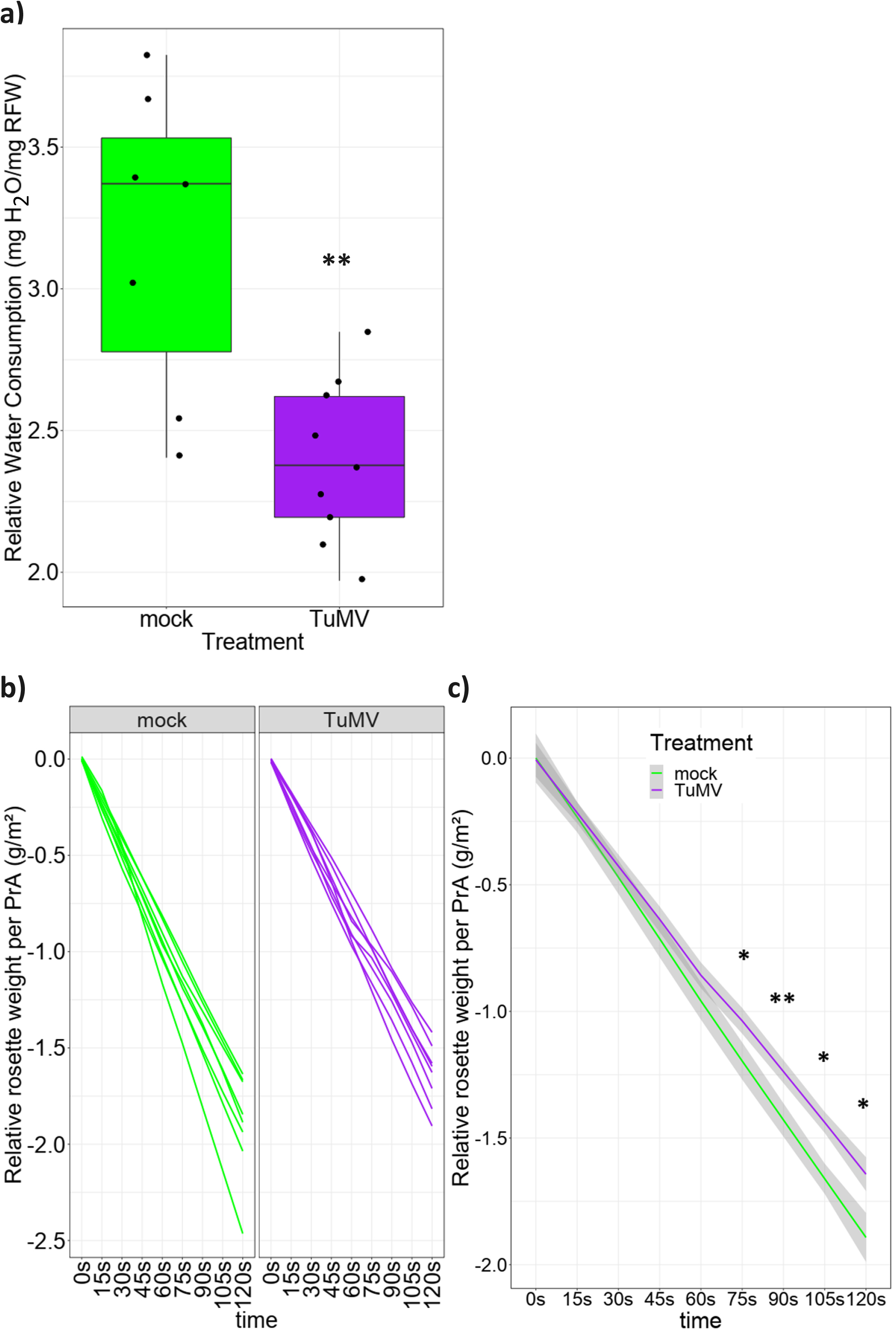
TuMV infection reduces relative water consumption. ***(a)*** Daily water consumption in TuMV-infected and mock-inoculated intact plants (12-13 DPI). RFW = Rosette Fresh Weight. Two independent experiments showing similar results were performed; combined results of both experiments are shown. N = 4-6 for each treatment and group. ***(b-c)*** Initial dehydration rate experiments. Rosette weight loss recorded through time per rosette projected area. Two independent experiments were performed showing similar results. Results from one representative experiment are shown, n = 8 for each treatment group. Spaghetti plots show ***(b)*** all rosettes separately and ***(c)*** smoothed conditional means by treatment (confidence intervals (95%) are shadowed in grey). Asterisks indicate statistically significant differences along time points.

### TuMV increases SA and ABA accumulation

Stomatal conductance is largely controlled by ABA and altered leaf water loss is often indicative of changes in accumulation or sensitivity to ABA (Verslues *et al.*, 2006). Also SA and JA are defense hormones that play a role as regulators of stomatal closure (Melotto *et al.*, 2017). Therefore, we quantified the level of these hormones in Arabidopsis rosette systemic tissue in infected and healthy plants (Figure 4). Under TuMV infection, both ABA and SA levels were greatly increased (205% and 404%, respectively; average of two independent experiments) whilst JA levels were not altered by TuMV infection.

**Figure 4.**
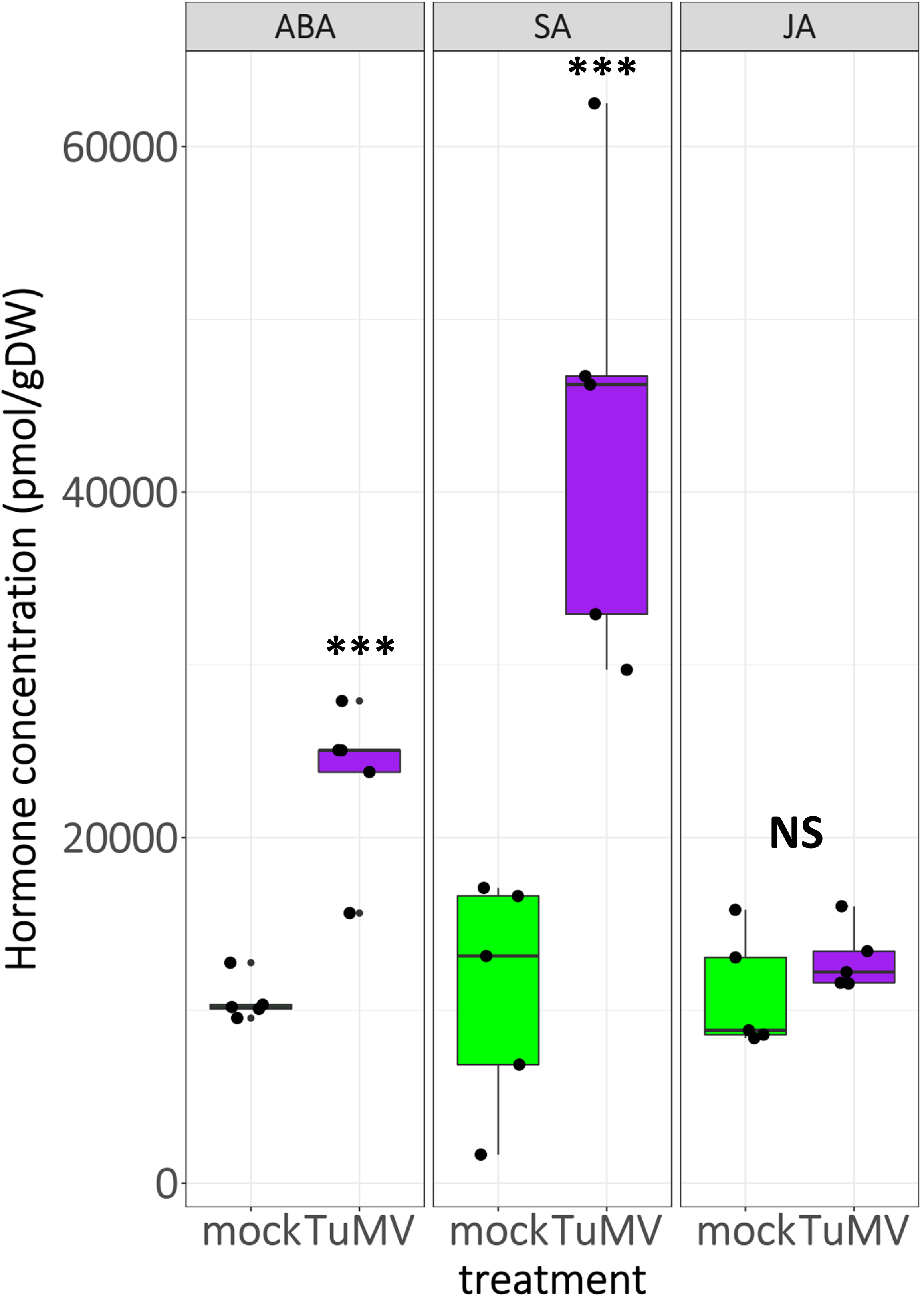
TuMV infection increased ABA and SA concentration. Accumulation of ABA, SA and JA were measured in the rosette of control and TuMV-infected plants. N = 5 per treatment group. Data from two independent experiments showed similar results. Results from one representative experiment is shown.

### ABA-insensitive mutant *abi2-1* prevent stomatal closure triggered by TuMV

Higher concentrations of defense hormones under TuMV infection were somewhat expected because higher levels not only of SA but particularly of ABA are traditionally regarded as underlying stomatal closure (Vishwakarma *et al.*, 2017; Ma *et al.*, 2018; Melotto *et al.*, 2017). To test for the effect of homeostasis of these hormones on the stomatal response to TuMV, we screened a panel of mutants in search of mutations which could abolish the TuMV-induced stomatal closure phenotype (Figure S4). We used *g_s_* measurements as a proxy for putative phenotype reversal. Neither mutants in production (*sid2*), perception (*npr1*) or degradation (*NahG*) of SA, nor the *aos* mutant (which is inactivated in the biosynthesis of JA and its derivatives), nor *lox1-1* (a mutant impaired in the JA-independent oxylipin pathway of stomatal closure response (Montillet *et al.*, 2013)), nor ABA-deficient mutant *aba3-1*, a severe mutant with only 30-50% normal ABA levels (Merilo *et al.*, 2017), could revert the phenotype. ABA signaling is also critical in the outcome of the physiological response triggered by this hormone. ABI2 (and other PP2CAs phosphatases) is repressed by ABA, de-repressing in turn OST1 (and other kinases of the SnRK family) to activate SLAC1 anion channels, promoting thus stomatal closure. PP2CA dominant positive/hypermorphic mutants *abi2-1* (Koornneef *et al.*, 1984; Leung *et al.*, 1997) cannot be repressed by ABA (Vlad *et al.*, 2009). Therefore OST1 is permanently repressed determining a high degree of stomatal aperture that is insensitive to exogenous application of ABA (Merlot *et al.*, 2002; Merlot *et al.*, 2001). Using the ABA-insensitive *abi2-1* signaling hypermorphic mutant the stomatal closure phenotype induced by TuMV was abolished (Figure 5). Moreover, a reproducible but marginally non-significant increase in stomatal aperture (Figure 5b) was detected under TuMV infection of *abi2-1* (18% higher mean stomatal aperture in infected plants and p = 0.076 for the combined analysis of two independent experiments). Therefore, our results suggest that JA and SA do not participate in TuMV-induced stomatal closure, and that reduced basal ABA levels (such as those present in *aba3-1* mutants) are compatible with this phenotype, whereas ABA downstream signaling is essential for it. This finding highlighted the relevance of ABA in stomatal closure mediated by TuMV but posed questions about how ABA homeostasis is modified in TuMV-infected plants resulting in decreased stomatal aperture.

**Figure 5.**
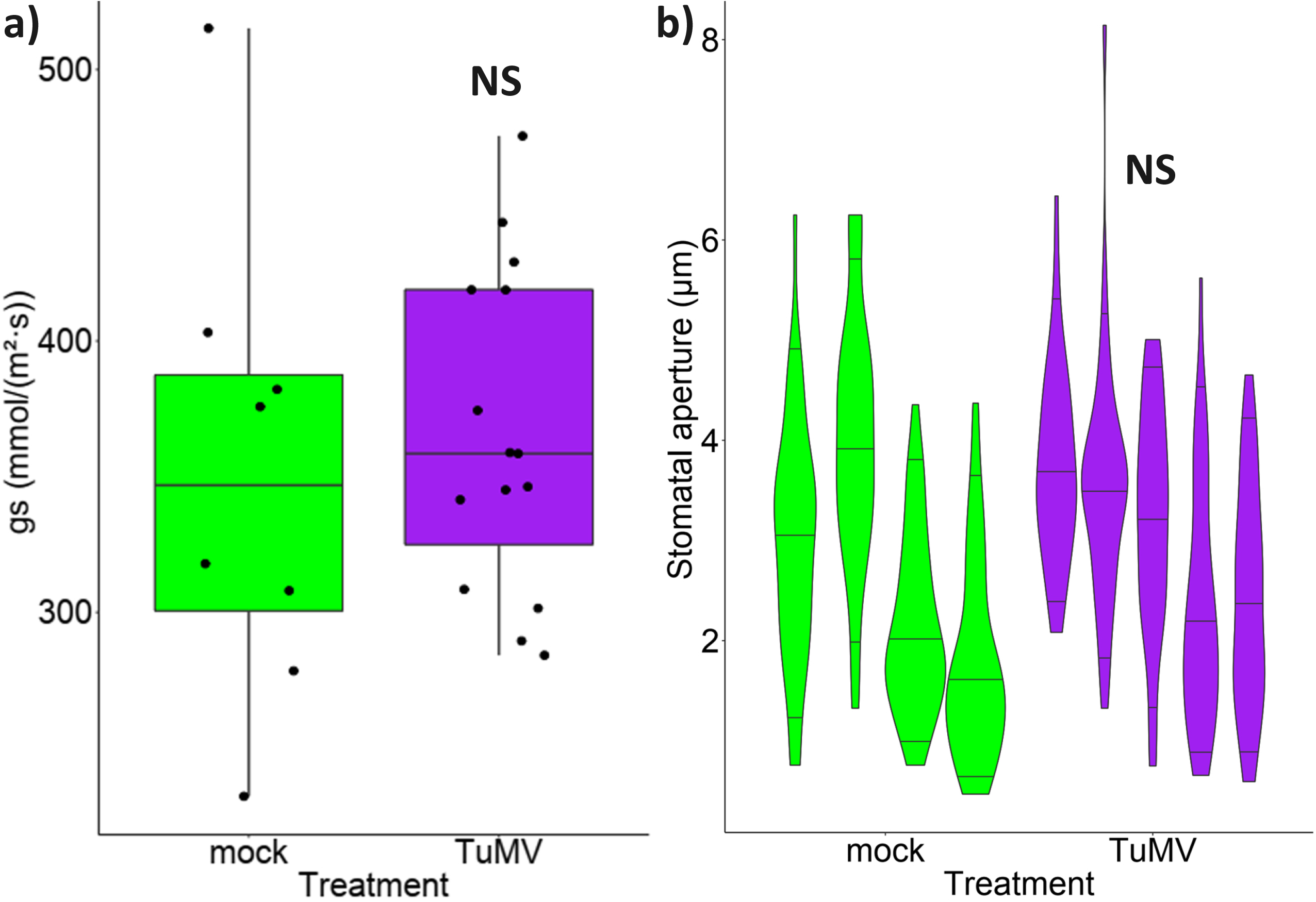
The ABA-insensitive *abi2-1* mutant abolishes TuMV-induced stomatal closure and exhibits a slight phenotype reversion. ***(a) G_s_*** in *abi2-1* control and TuMV-infected plants at 12 DPI. N = 8 for mock and n = 16 for TuMV. ***(b)*** Stomatal aperture in *abi2-1* control and TuMV-infected plants at 12 DPI. Violin plots show 0.05, 0.5 and 0.95 quantiles for stomatal aperture data (n = 40) from each individual. Data from two independent experiments showed similar results. Results from one representative experiment is shown, n = 5 for each treatment group, p > 0.05. Ler control plants showed the same *g*_*s*_ response to TuMV infection as Col-0 (data not shown).

### TuMV alters ABA production, catabolism, signaling and transport

ABA homeostasis during plant stress responses is modulated by its production, inactivation, and transport and is considered to be achieved mainly through transcriptional regulation (Ma *et al.*, 2018; Kushiro *et al.*, 2004). Therefore, we investigated which step(s) of ABA homeostasis is responsible for its over-accumulation under TuMV infection. As ABA metabolism occurs both in shoot and root tissues (Park *et al.*, 2017; Kushiro *et al.*, 2004), aerial and root parts from hydroponically-grown plants were separately harvested to quantify the expression of the most relevant genes related to ABA biosynthesis, degradation, signaling and transport (Figure 6). ABA biosynthetic genes are analyzed in Figure 6a. The rate-limiting step in ABA biosynthesis is catalyzed by the plastid-encoded *NCED3* gene (Vishwakarma *et al.*, 2017; Ma *et al.*, 2018), which is the major stress-induced NCED in leaves and, together with NCED2, account for the NCED activity in roots (Tan *et al.*, 2003). Upon TuMV infection, NCED3 was down-regulated in shoots. ABA2 acts downstream of NCED3 in the cytoplasm (Ma *et al.*, 2018) and was also down-regulated by TuMV. Contrarily, AAO3, which catalyzes the last step in ABA biosynthesis (Ma *et al.*, 2018), was up-regulated by TuMV. In roots, NCED3 and ABA2 were not altered by TuMV infection and AAO3 showed a level of up-regulation similar to that of shoots. Catabolism plays an important role in ABA homeostasis and a fine-tuned balance of transcriptional activity exists between biosynthetic and degradation genes during stress and recovery (Kushiro *et al.*, 2004). The Arabidopsis P450 genes of the CYP707A family CYP707A1–CYP707A4, encode ABA 8’-hydroxylases that catabolize ABA (Ma *et al.*, 2018; Kushiro *et al.*, 2004). Of these, CYP707A1 and A3 are important mainly in root tissue whereas CYP707A3 is more prominent in leaves (Kushiro *et al.*, 2004). CYP707A1 and A3 were both down-regulated in root and shoot tissues (Figure 6b). Therefore, higher endogenous ABA levels under TuMV infection appear to be due to reduced catabolism compensating for reduced biosynthesis (Figure 6a,b). ABI2 plays a central role acting as a negative regulator of ABA signaling (Merlot *et al.*, 2001; Lim *et al.*, 2015). In addition to its well-known role in stomatal closure, ABI2 was also reported to have a role in root growth (Leung *et al.*, 1997). We investigated in public gene expression databases ABI2 expression patterns regarding anatomy and perturbations or stresses. An inspection using the eFP-Browser tool (Winter *et al.*, 2007) of ThaleMine (Krishnakumar *et al.*, 2015) and Genevestigator V3 (Hruz *et al.*, 2008) showed that ABI2 is mainly expressed in guard cells and roots and is induced by ABA, salt and drought, and senescence (Figure S5), the latter being a process that is greatly accelerated by TuMV-UK1 (Manacorda *et al.*, 2013). Accordingly, we observed an increase in ABI2 expression in roots as well as a weak up-regulation in shoots (Figure 6c).

**Figure 6.**
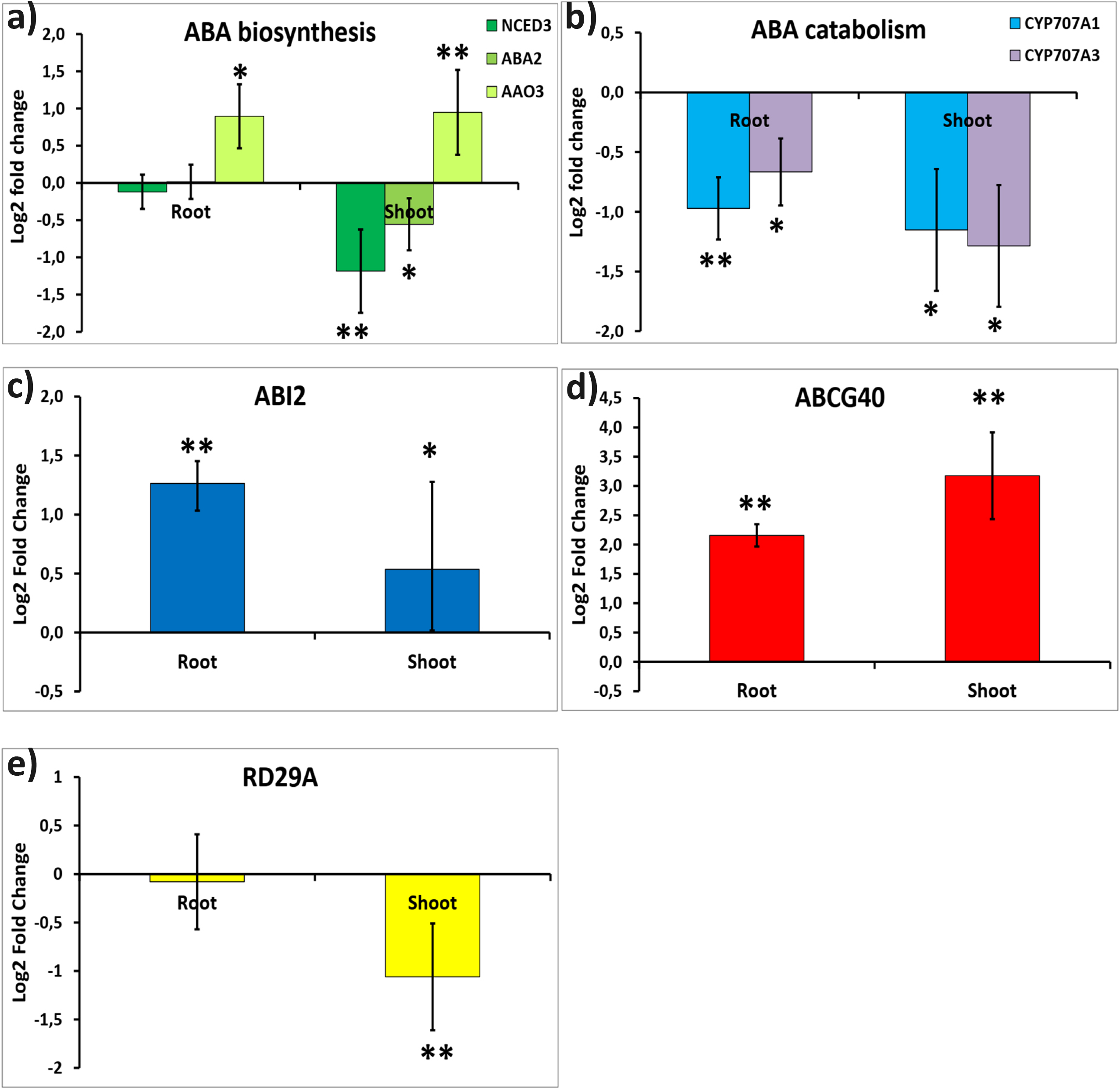
TuMV infection alters relative transcript accumulation of key genes in ABA metabolism and function. QPCR analyses of genes involved in ABA biosynthesis ***(a),*** catabolism ***(b)***, signaling ***(c)***, transport ***(d),*** and stress response ***(e)***. Bars = SE. N ≥ 4 per group.

Besides higher overall rosette ABA concentration triggered by TuMV (Figure 4), augmented ABA transport into guard cells could also be responsible of stomatal closure under infection. ABCG40 is a plasma membrane-bound ABA transporter mainly expressed in guard cells of aerial tissues but also in primary and lateral roots (Kang *et al.*, 2010; Kuromori *et al.*, 2018). It has very high affinity for ABA and imports it into the guard cells from the surrounding tissues (Park *et al.*, 2017; Ma *et al.*, 2018) where ABA participates in stomatal closure (Kang *et al.*, 2010). To gain insight into ABA transport processes under TuMV infection, we measured ABCG40 expression in roots and shoots. ABCG40 was highly over-expressed in roots (4.5X-linear fold) and shoots (9X-linear fold) (Figure 6d). The dehydration and pathogen responsive gene RD29A was downregulated by TuMV in shoots (Figure 6e). This result is consistent with RD29A expression reported as downregulated by TuMV and several other biotic stresses including another virus infecting *Brassicaceae* (Table S4). The results of these qPCR experiments are summarized in Table S4. They are compared with the reported response of these genes to main stresses (perturbations) and tissue and stage of development expression pattern as informed by the online databases of Genevestigator. Table S4 shows that under TuMV infection, the expression of the selected genes for ABA homeostasis and stress response analyzed here, coincide to a great extent with their reported responses to ABA, including abiotic stresses that typically induce ABA responses, and also to their responses to biotic stresses.

### TuMV increases drought susceptibility in Arabidopsis

Stomatal closure and reduced leaf water loss are well studied phenomena that lead to improved drought tolerance (Lim et al., 2015; Verslues et al., 2006). The overexpression of ABCG40 (Figure 6d), together with stomatal closure (Figure 2) and reduced leaf water loss (Figure 1, Figure S3, Figure 3) all appeared to point out to that outcome under TuMV infection. However, a previous report showed that TuMV increased susceptibility to water deficit in Arabidopsis (Prasch and Sonnewald, 2013). Hence, we performed experiments combining drought and TuMV infection. Soil drying is a technique often used to evaluate plant response to water deficit. In the literature, two main approaches are found: in one of them one plant is used per pot and a treatment/genotype is randomly assigned to each one (Westwood *et al.*, 2013; Xu *et al.*, 2008; Prasch and Sonnewald, 2013), while in the other several plants are placed in the same pot and different treatments are assigned to the plants (Xu *et al.*, 2008). This latter approach ensures that roots of plants receiving different treatments, experience the same water deficit stress (Verslues *et al.*, 2006). In particular, plants with decreased stomatal conductance or decreased growth and leaf area (phenotypes which are all present in TuMV-infected plants) are expected to deplete soil water more slowly to avoid low water potential, and may be erroneously designated as more tolerant than controls (Verslues *et al.*, 2006). To determine whether this is indeed the case in our experimental conditions, we firstly tested the one-plant-per-pot approach (Figure S6). It can clearly be seen from Figure S6 that mock-inoculated plants consume available water at a much faster rate than their TuMV-infected counterparts. At the end of the experiment at 42 days after water withholding (DAWW), pots with mock-inoculated plants are still alive and had started reaching a weight plateau, indicative of complete available water depletion in the pot. Meanwhile, some of the TuMV-infected plants had already died because of the advanced stage of viral infection (Figure S6a). This is in line with the more susceptible phenotype detected by (Prasch and Sonnewald, 2013), who used separate pots per plant. Notwithstanding, pots containing TuMV-infected plants had still around 20 g more of available water. Therefore, we carried out experiments with the two-plant-per-pot approach, assuring that plants receiving different treatments experienced the same water deficit stress (Figure 7). Given the presence of two plants per pot, water was more rapidly removed from soil and at 26 DPI = DAWW, plants receiving each treatment were severely stressed, showing evident leaf turgor loss (Figure 7a). After re-watering however, 100% of mock-inoculated plants survived but only 9% of TuMV-infected plants recovered (Figure 7b; χ^2^ = 7.3e-11, Table 1). To test for the robustness of the phenomenon and to facilitate the comparison with experimental conditions reported for the combined stress of drought + TuMV (Prasch and Sonnewald, 2013), we repeated the experiment but in LD conditions and compared pot weight loss and survival rates under different photoperiod conditions and treatments (Figure 7c-d, Table 1). As expected, soil drying was much faster in LD conditions (Figure 7c-d), to the extent that this standard photoperiod is perhaps not the most suitable for our pathosystem to study in detail the low water potential response (Verslues *et al.*, 2006). Nonetheless, we found that TuMV increased drought susceptibility also in LD experiments (χ^2^ = 0.02, Table 1). These results are in line with reduced expression of RD29A gene in systemic tissues from well-watered TuMV-infected plants in both SD and LD conditions (Figure 6e, Figure 7e). Of note, previous results (Manacorda *et al.*, 2013) studying TuMV infection in well-watered Arabidopsis under similar LD growing conditions, allowed plants to survive for more than 21 DPI, indicating that TuMV alone is not responsible for premature plant death observed here. Therefore, the two-plants-per-pot approach, which is appropriate to study the combined stresses of TuMV infection + drought, showed that despite stomatal closure and reduced relative evapo-transpiration and water consumption, TuMV-infected plants are more susceptible to drought than mock-inoculated controls. The observed reduction in RD29A expression under well-watered conditions, both in SD and LD photoperiods, is in line with the reported expression of this gene under pathogen attack and SA treatments (when it is down-regulated, Table S4) and contrary to its expression under water stress only (when it is up-regulated (Narusaka *et al.*, 2003)). We confirmed that SA treatment alone downregulates RD29A (Figure S7).

**Table 1.** Chi-square tests for LD and SD experiments showing reduced Arabidopsis tolerance to drought under TuMV infection. One representative experiment is shown.

| Condition | | Mock | TuMV | Adjusted residuals: | $\chi^2$ test (Fisher's exact) |
| --- | --- | --- | --- | --- | --- |
| LD | Alive | 23 | 17 | 2,6268 | **0,021554** |
| LD | Dead | 0 | 6 |  |  |
| SD | Alive | 23 | 2 | 6,2161 | **7,2874E-11** |
| SD | Dead | 0 | 21 |  |  |

**Figure 7.**
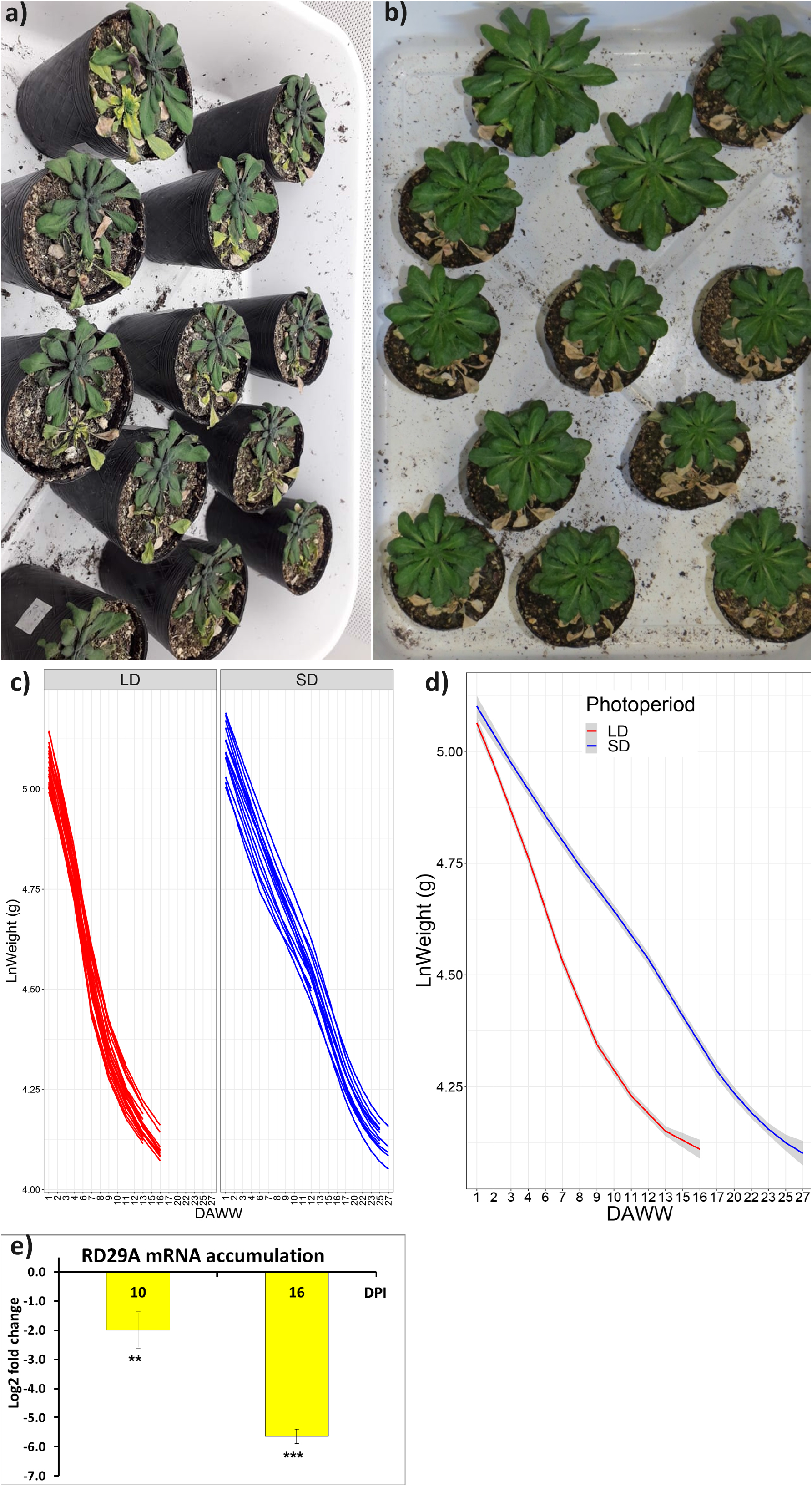
TuMV infection increases drought susceptibility in Arabidopsis. ***(a-b)*** Images showing plants at 26 DPI and DAWW and just prior re-watering ***(a)*** and 5 days after re-watering ***(b)***. Plant were grown in short day conditions. Mock-inoculated plants are placed at top for each pot, TuMV-infected plants next to them. ***(c-d)*** Comparison between drought experiments in short day (SD) and long day (LD) conditions. Line graphs show the kinetics of water loss (LnWeight) for pots containing two plants each, either in SD or LD conditions. Line graphs show each pot ***(c)*** separately and ***(d)*** smoothed conditional means by treatment (confidence intervals (95%) are shadowed in grey). Two independent experiments rendered similar results. Results of one representative experiment are shown. ***(e)*** RD29A relative expression under TuMV infection in well watered LD conditions.

### ORMV also triggers a hot-leaf phenotype in Arabidopsis

T_leaf_ increase is not a universal phenomenon arising from compatible viral infections. For example, tobacco plants susceptible to TMV (a virus belonging to the *Tobamoviridae* family unrelated to TuMV) failed to accumulate SA and did not increase T_leaf_ relative to healthy controls (Chaerle *et al.*, 1999). Therefore, we tested whether Arabidopsis infected with ORMV (a *Tobamoviridae* member capable of establishing a compatible infection in this host) showed a hot-leaf phenotype. Figure S8 shows that this is indeed the case.

## Discussion

### Plant viruses and its impact on stomata

TuMV induced an average T_leaf_ increase of 0.5-1 °C (Figures S1-S2 and Figure 1), which is within the reported range for other virus-infected plants (Aguilar *et al.*, 2017; Chaerle *et al.*, 2006). As expected, TuMV decreased *g_s_* (Figures 1d and Figure S3) in line with reduced stomatal aperture (Figure 2) and lowered relative daily water loss and initial dehydration rate (Figure 3). This characterization of the stomatal response of Arabidopsis to TuMV adds another piece of information about this well-studied pathosystem.

High relative T_leaf_ profile or/and diminished *g_s_* or shoot water loss under viral infection have been reported previously for other pathosystems (Aguilar *et al.*, 2017; Xu *et al.*, 2008; Murray *et al.*, 2016). This is, however, not a universal phenomenon; in the case of Arabidopsis, there are reports describing enhanced water loss and/or decreased T_leaf_ in virus-infected plants (Westwood *et al.*, 2013; Aguilar *et al.*, 2017) concomitantly with enhanced drought tolerance, a pattern opposite to what we have found for the Arabidopsis-TuMV interaction (see below for further discussion). Our findings are however in line with those of (Guo *et al.*, 2005), who detected a sharp decrease in *g_s_* in TuMV-infected *Brassica juncea* plants (a member of the *Brassicaceae* family related to Arabidopsis). We also found an increase in T_leaf_ in ORMV-infected Arabidopsis plants (Figure S8), whereas (Chaerle *et al.*, 1999) did not find any T_leaf_ differences when infecting tobacco plants with TMV, a virus related to ORMV. These contrasting findings highlight the need to study in detail each plant-virus interaction to characterize the impact that each virus may have on stomatal function.

### TuMV increased SA and ABA accumulation but stomatal closure depends on ABA perception and not wild-type basal levels

ABA, SA and JA are plant hormones involved in antiviral defense (Klessig *et al.*, 2018; Vlot *et al.*, 2009; Alazem and Lin, 2015; Alazem and Lin, 2017; Lewsey and Palukaitis, 2010) and drought response (Wang *et al.*, 2018; Raskin, 1992; Okuma *et al.*, 2014) and a recent and growing corpus of research is making links between these two roles (Li *et al.*, 2012; Aguilar *et al.*, 2017). Interestingly, a synergistic interaction between ABA and SA was proposed for at least some aspects of antiviral responses (Alazem and Lin, 2015). JA levels were mostly unchanged in response to TuMV infection (Figure 4), a result previously found in this pathosystem (Casteel *et al.*, 2015). On the other hand, both ABA and SA levels were increased (Figure 4). SA synthesis/signaling and the SA-dependent defense are important to limit viral infection and spread (Rodriguez *et al.*, 2014; Conti *et al.*, 2012; Collum and Culver, 2016). However, the role played by SA in the Arabidopsis-TuMV interaction is not yet fully understood: on the one hand the use of transgenic plants overexpressing *NahG* with reduced levels of SA (Brenya *et al.*, 2016) led to the conclusion that endogenous SA did not affect resistance to TuMV (at least from the point of view of visual symptom development). On the other hand, previous studies using similar *NahG* transgenic plants found that SA plays a role in ROS generation after TuMV infection (Manacorda *et al.*, 2013). The same study found that a set of SA-responsive genes related to defense and senescence was highly induced from early stages of TuMV infection with the UK1 strain. Additionally, (Poque *et al.*, 2018) showed that the interaction between TuMV protein HC-Pro and a host SA-binding protein reduced SA signaling capability and increased susceptibility to TuMV, suggesting that the SA pathway is an important obstacle to TuMV spread. Our findings here are in line with previous studies that showed an increase in SA levels after TuMV infection in Arabidopsis (Poque *et al.*, 2018; Casteel *et al.*, 2015). However, increased levels of SA are unlikely to be behind the stomatal closure triggered by TuMV, as mutations affecting SA biosynthesis or perception, or decreasing of SA levels through *NahG* overexpression failed to prevent a reduction in *g_s_* (Figure S4). Perhaps less unexpectedly due to unchanged JA levels in response to TuMV infection, oxylipins pathways mutants (either related with the JA biosynthetic pathway-*aos*- or not -*lox1-1*) also failed to abolish the phenotype triggered by TuMV. Interestingly, *aba3-1* plants also failed to suppress the TuMV- induced decrease in *g_s_*, demonstrating that reduced endogenous basal ABA levels alone are not enough to hinder this response. The fact that *aba3-1* mutants are still capable of closing stomata despite their low endogenous ABA levels in an ABA-dependent process could be explained on the basis that not only the minimum ABA threshold necessary to trigger stomatal closure is not yet determined, but also because it is proposed that the threshold of ABA to initiate downstream processes is lower in the guard cells (Merilo *et al.*, 2017). ABA-insensitive dominant positive/hypermorphic *abi2-1* mutant plants abolished stomatal closure and low *g_s_* under TuMV infection (Figure 5), proving that these phenotypes are dependent on ABA in general and ABA sensitivity in particular. In WT plants, ABI2 represses OST1/SnRK2.6, which in turn acts activating SLAC1 anion channels that trigger stomatal closure in guard cells (Lee *et al.*, 2013). In *abi2-1* mutants, OST1 is permanently repressed determining a high degree of stomatal aperture (Merlot *et al.*, 2002; Merlot *et al.*, 2001). Moreover, stomatal aperture is further promoted in these mutants because PP2CA proteins can also directly repress SLAC1 anion channels (Rodriguez *et al.*, 2019). To sum up, TuMV-triggered stomatal closure is dependent on ABA signaling by PP2CAs (Figure 5) but not on other well-known hormones or metabolites capable of causing stomatal closure and this phenotype does not require WT basal rosette ABA levels despite occurring concomitantly with a rise in endogenous ABA levels (Figure 4).

### TuMV impacts on ABA metabolism and transport

The understanding of the important role that ABA response has in the context of viral infection is gaining momentum (Alazem and Lin, 2017; Pasin *et al.*, 2020). For example, ABA has been proposed to be at the core of extreme resistance mechanisms following soybean infection by the potyvirus Soybean Mosaic Virus (SMV) (Alazem *et al.*, 2018; Alazem, Widyasari, *et al.*, 2019). Here, we performed transcriptomic qPCR experiments to understand the origin of increased overall rosette ABA levels under infection and its putative effects on pathogen response as well as the relative independence of stomatal closure on absolute endogenous ABA levels. Increased levels of ABA in shoots were detected in spite of NCED3 and ABA2 down-regulation in shoots (Figure 6a). The slight relative up-regulation of AAO3 in roots and shoots could account for a faster processing of remaining ABA-aldehyde intermediate, but this conclusion is not guaranteed since it has been shown that AAO3 mRNA could be strongly induced without downstream higher protein level or enzymatic activity (Ma *et al.*, 2018). Interestingly, AAO3 (but not NCED3 or ABA2) induction was also reported in Arabidopsis infected with BaMV (Alazem *et al.*, 2014). NCED3 and ABA2 were found to be positive regulators of BaMV accumulation and AAO3 and several ABA signaling downstream of it (including the hypermorphic *abi1-1* that is functionally equivalent to *abi2-1* used here) had the inverse role. Therefore, NCED3 and ABA2 down-regulation and AAO3 and ABI2 overaccumulation detected in our study (Figure 6) are most probably part of a defense response mounted by the host. (Alazem *et al.*, 2014) found that both CMV and BaMV triggered ABA accumulation but also upregulated the mRNA levels of several genes from the SA and ABA pathways, therefore concluding that both CMV and BaMV infection disrupt ABA-SA antagonism. Our results support their suggestion that the increase of ABA and upregulation of the ABA and SA pathways may be common features of RNA viruses infection (Alazem *et al.*, 2014). Additionally, down-regulation of P450 cytochromes that degrade ABA (Figure 6b) could be compensating for the reduced biosynthesis, also taking into account that in roots NCED3 and ABA2 have unchanged transcriptional levels but P450 cytochromes are also downregulated there and ABA transport throughout the plant is a well-established phenomenon. Further studies should complement these findings by analyzing ABA content in shoots and roots separately. This molecular phenotype is in agreement with the augmented drought susceptibility detected (Figure 7) since it was recently suggested that drought-induced ABA synthesis occurs in shoot vasculature (Park *et al.*, 2017) and in Arabidopsis, NCED3 is considered as a dominating contributor to ABA production under water deficit, with other NCEDs, as well as other enzymes within ABA synthesis pathway playing a relatively minor role (Ma *et al.*, 2018). NCED gene was found to be overexpressed in response to drought stress in several plant species including Arabidopsis (Vishwakarma *et al.*, 2017). Therefore, NCED3 down-regulation by TuMV goes in the opposite direction of water stress tolerance. NCED3 is a chloroplastic gene and its transcription downregulation could originate from its subcellular localization and the early onset of senescence triggered by TuMV (Manacorda *et al.*, 2013). Recently, more attention was given to chloroplast function impairment during viral infections, as well as the role this organelle has in antiviral defense (Zhao *et al.*, 2016; Bhattacharyya and Chakraborty, 2018). Transcriptomic studies of potyviral infections found that plastid genes and genes involved in chloroplast functions were preferentially underexpressed, whereas senescence-associated genes (SAGs) including several defense- and stress-associated genes, were overexpressed (Revers and García, 2015). Moreover, the majority of Potyviral proteins have the capability of physically interacting with host chloroplastic proteins (Revers and García, 2015) and TuMV can alter chloroplast morphology (Zhao *et al.*, 2016). TuMV UK1 strongly induces the SAG ORE1 transcription and early chlorophyll loss in systemic Arabidopsis and *B. juncea* leaves (Manacorda *et al.*, 2013). ORE1 is a negative regulator of G2-like transcription factors, which are important in chloroplast development and maintenance (Rauf *et al.*, 2013). In line with this, it was shown that TuMV-infected *B. juncea* plants showed a noticeable decrease in total chlorophyll content after 2 weeks of infection (Guo *et al.*, 2005).

Not only overall ABA levels but also ABA distribution within the plant is important to trigger physiological responses. ABCG40 is the only *bona fide* ABA importer into guard cells reported to date (Park *et al.*, 2017; Kuromori *et al.*, 2018). Contrary to ABA exporters from guard cells like ABCG25, in shoots ABCG40 is mostly expressed in guard cells (Kuromori *et al.*, 2010; Kuromori *et al.*, 2018; Kang *et al.*, 2010). Although ABA can be synthesized in guard cells to allow for rapid stomatal responses to atmospheric factors, long-term stomatal stress responses appear to depend more on ABA biosynthesized in phloem companion cells (Merilo *et al.*, 2015; Kuromori *et al.*, 2018). Therefore, increased levels of ABCG40 mRNA (Figure 6d) could contribute a major part of the ABA pool in guard cells. Additionally, although contributing to a minor part to the ABA pool in shoots, CYP707A1 can down-regulate ABA content in guard cells via its preferential expression there, thus influencing stomatal closure (Kuromori *et al.*, 2018). CYP707A1 and CYP707A3 were down-regulated both in roots and shoots (Figure 6b). Accordingly, increased ABA transport to guard cells and reduced catabolism could explain stomatal behavior under TuMV infection even in *aba3-1* mutants with lower overall shoot ABA levels (Figure S4). Down-regulation of CYP707A genes were unexpected in the light of the augmented ABA concentration detected in TuMV-infected rosettes (Figure 4) because CYP707A genes are induced by high endogenous ABA, whose levels contribute to limit under stress (Kushiro *et al.*, 2004). CYP707As down-regulation detected here are also the contrary pattern of what is generally detected in plants under water stress (Ma *et al.*, 2018; Kushiro *et al.*, 2004), where these cytochromes control the excessive level of ABA, further indicating that TuMV-infected plants are ill-prepared to stand drought. PP2CAs including ABI1 and ABI2 are required for ABA signaling in Arabidopsis and generally function as negative regulators of the ABA response (Lee *et al.*, 2013; Merlot *et al.*, 2001). ABI2 has been reported as transcriptionally up-regulated by ABA and abiotic stresses (Figure S5c-d, Table S4), and regarding pathogen response, at least some PP2CA members are important in determining bacterial resistance via stomatal closure through their interaction with ABA receptors (Lim and Lee, 2015). ABI2 mRNA relative levels were marginally higher in TuMV-infected shoots (Figure 6c). This effect, in fact, could be under estimated by the whole-shoot RNA extraction, that reflects the effect of overall shoot ABA levels. Since in shoots guard cells are the primary site of ABI2 expression (Figure S5a), higher ABA levels determined by increased ABCG40 and decreased CYP707As transcription (Figure 6b,d) were expected to determine high local mRNA ABI2 levels (Figure S5c). The strong overexpression of the high-affinity importer ABCG40 in shoots (+9.5 fold relative to control plants) suggests that ABA is being massively imported from surrounding tissues into guard cells. Nevertheless, higher ABA levels in guard cells must lead to increased local ABI2 protein repression: ABA perception through PYR/PYL/RCAR receptors leads to interaction with and inactivation of PP2CAs like ABI2 (Rodriguez *et al.*, 2019), thus promoting the detected stomatal closure (Figure 2). In this context, it is interesting to note that stomatal aperture assays on *abi2-1* hypermorphic mutants (Figure 5b) although showed no statistical differences between treatments, yielded similar results in two independent experiments, with an average stomatal aperture increase of 18% in TuMV-infected plants instead of a reduction as detected in infected WT plants (Figure 2). Accordingly *g_s_* measurements, carried on in plants of additional independent experiments (Figure 5a), was detected to be slightly higher in TuMV-infected *vs*. healthy *abi2-1* mutants, an opposite pattern regarding WT results (Figure S3, Figure 3). We propose that in these mutants, the local increase in ABA in guard cells under infection leads to an expected increase in ABI2 mRNA, which is translated into additional mutant ABI2 protein, which is in turn incapable of down-regulation through PYR/PYL/RCAR ABA receptors, further promoting stomatal aperture.

### Transcriptomic profile of ABA metabolic pathway genes under TuMV marked induced senescence, pathogen response and increased drought susceptibility

Table S4 compares our transcriptomic results for the selected ABA metabolic pathway genes with reported data retrieved from Genevestigator V3 (Hruz *et al.*, 2008) for most important perturbations (stresses), tissues/cell lines (anatomy) and developmental expression pattern in Arabidopsis. Except for RD29A, the expression pattern of all genes analyzed under TuMV infection was the same as reported for senescent tissues (Table S4, Figure 6). This was expected since it has been reported that TuMV induces the early onset of senescence in Arabidopsis (Manacorda *et al.*, 2013; Sánchez *et al.*, 2015). The majority of the genes analyzed here (Figure 6) are markedly altered by biotic stresses, particularly biotrophic pathogens that typically trigger a SA-dependent response (Li *et al.*, 2019) like *P. syringae*, but also *P. parasitica* and *M. incognita*, in the same direction of regulation as triggered by TuMV (up-or down-regulation). Most of these genes were also highly expressed in roots and guard cells, as expected, as they are places where ABA plays important physiological roles. Interestingly, ABCG40 expression had been previously detected in roots and its expression increased mainly under pathogen attack, as it did here (Figure 6d). As for RD29A expression, its consistent down-regulation by TuMV both under SD and LD conditions (Figure 6e, Figure 7e) shows a pattern that is the opposite of what was reported under drought conditions (Wang *et al.*, 2018; Yamaguchi-Shinozaki and Shinozaki, 1994).

### RD29A is downregulated by SA and viruses

RD29A is routinely used as an indicator of stress response, since it was originally reported to be induced by ABA treatment but also by abiotic stresses such as salt and drought, for which promoter elements exist that can respond independently from ABA (Yamaguchi-Shinozaki and Shinozaki, 1994). Interestingly, (Westwood *et al.*, 2013) confirmed the up-regulation of RD29A gene under ABA treatment but also found that RD29A was repressed in Arabidopsis plants infected with Fny-CMV in both well-watered and drought-stressed conditions. These authors found no changes in ABA levels following infection. Remarkably, data mining using Genevestigator revealed that previous reports showed that viral infections triggering an SA response repress RD29A both in the case of TuMV (Yang *et al.*, 2007) and CaLCuV (Ascencio-Ibáñez *et al.*, 2008) and that SA treatment alone represses RD29A in plantlets and mature rosettes (Table S4), a result we confirmed here (Figure S7). Noteworthy, the strength of RD29A repression appear to be linked to the duration of the SA treatment: when a single SA spray is used, RD29A is significantly but slightly repressed at 3-6 h post treatment but this effect fades away at 24 h or later (Figure S7, Table S4). On the contrary, a sustained SA treatment such as a 1-day submersion of Arabidopsis roots in SA solution triggers a strong repression of RD29A (−13.6 X linear fold change, Table S4). This kind of sustained response is more likely to be occurring in our TuMV-infected plants at 12 DPI, when infected plants have a near 400% increase of SA levels (Figure 4). Indeed, previous reports showed that a transcriptomic and biochemical systemic response that is typically SA-dependent is triggered by TuMV (UK1) as early as 4 DPI, through 21 DPI (Manacorda *et al.*, 2013). The fact that TuMV-induced RD29A repression in LD conditions is severely accentuated from 10 to 16 DPI, further supports that view (Figure 7e). Moreover (Ascencio-Ibáñez *et al.*, 2008), showed that CaLCuV (a virus that triggers a pathogen response via the salicylic acid pathway) highly repressed RD29A at 12 DPI under SD conditions. Additionally, previous reports using *npr1*, *sni1* and *brca2a* single, double or triple mutants (Durrant *et al.*, 2007; Wang *et al.*, 2010) demonstrated that RD29A depends on these SA-responsive genes for its normal transcriptional levels. The SA response of RD29A was even reversed in some of these mutant’s combinations. Therefore, RD29A appears to be not only an indicator of ABA-mediated abiotic stress response (when it is overexpressed) but also of biotic stress and particularly, viral infections and SA treatment (when it is underexpressed) (Table S4, (Westwood *et al.*, 2013; Ascencio-Ibáñez *et al.*, 2008; Yang *et al.*, 2007), this study: Figure 6e, Figure 7e, Figure S7). This response is not bound to lower ABA levels ((Westwood *et al.*, 2013), this study: Figure 4) and could arise from the independently acting cis-elements described in its promoter region (Yamaguchi-Shinozaki and Shinozaki, 1994). Here, we detected both ABA and SA increased levels (Figure 4). These hormones of opposite effect on RD29A expression resulted in a pattern of expression that is not the same as reported after ABA increase only, but should not be unexpected: when confronted to a combination of stresses, plants mount unique physiological, molecular, and biochemical responses that is not just an additive effect of the responses deployed when the stresses are imposed separately (Mahalingam, 2015). In fact, regarding specifically dual stress effects on RD29A, a previous work studied RD29A expression under combined ABA + NaCl treatments and found that not only the magnitude of RD29A expression but also its temporal dynamics were altered relative to the response recorded after single stimulus, despite NaCl being known to trigger ABA itself (Lee *et al.*, 2016). Accordingly, despite enhanced ABA levels during TuMV infection, RD29A expression would be ultimately determined from the complex weighting between ABA and SA in the regulation of this gene. Plants that cannot induce RD29A in the presence of ABA are also more susceptible to NaCl or mannitol (Jeong *et al.*, 2018) and we found here that TuMV-infected plants were more susceptible to drought stress (Figure 7).

### Decreased drought tolerance under TuMV infection: putative physiological and molecular causes

TuMV induced stomatal closure (Figure 2) and reduced *g_s_* (Figure S3, Figure 3) concomitantly with reduced drought tolerance (Figure 7a,b, Table 1). These apparently contrasting phenomena could have their explanation in the broader role that SA and particularly ABA have in regulating water stress in plants and in the spatial distribution of ABA (Wang *et al.*, 2018; Kuromori *et al.*, 2018; Yoshida *et al.*, 2019). The strong induction of high-affinity ABCG40 (Figure 6d) under TuMV could indicate that an important fraction of additional ABA accumulated could be translocated to specific cell types such as guard cells, leaving less hormone available to other physiological functions that cope with stress. A recent review (Yoshida *et al.*, 2019) summarizing ABA roles under non-stress conditions specifically noted that the uneven distribution of ABA concentration throughout the leaf due to the action of transporters, catalytic enzymes and ABA mobility can explain stomatal closure under well-watered conditions and that this effect could be obscured by dilution in whole-leaf ABA measurements. The authors also pointed out that measuring ABA with cellular resolution remains an important and outstanding task for the future. Physiological ABA functions extend well beyond stomatal closure, coordinating all the various aspects of drought response (Verslues *et al.*, 2006) including water penetration and hydraulic control, both in leaves and roots to systemically cope with water stress (Kuromori *et al.*, 2018; Yoshida *et al.*, 2019). In roots, ABA regulates not only hydraulic conductance (possibly by regulating aquaporin activity (Kuromori *et al.*, 2018)) but also primary growth, suberization and hydrotropism (Yoshida *et al.*, 2019) and it can inhibit lateral root formation under NaCl stress (Duan *et al.*, 2013). The modification of these traits can obviously have an impact in water stress tolerance. Interestingly, the latter authors found that lateral root inhibition occurs in root endodermis cells, a cell line where NaCl and other stresses reportedly induced ABCG40 and ABI2 (Figure S5, Table S4), whose transcripts we detected highly overexpressed in roots under TuMV infection (Figure 6c,d). Noteworthy, both ABCG40 and ABI2 were reported to be highly expressed not only in guard cells, a feature that can explain that an over-expression would determine stomatal closure (Figure S3, Figure 3, Figure 6c,d, shoot analyses), but also in root endodermis and root cortex and lateral root cap, where PP2CAs like ABI2 were recently found to be involved in root patterning under stress (Rodriguez *et al.*, 2019; Dietrich *et al.*, 2017; Orman-Ligeza *et al.*, 2018).

Therefore, increased reported ABCG40 levels in roots occurs in cell lines that are highly responsive to ABA and leads to important morphological changes, pointing out to a possible additional role for this ABA transporter in root tissue besides its role in shoots (Kuromori *et al.*, 2018). As TuMV infection induced root overexpression of ABI2 and ABCG40 genes (Figure 6c,d), we speculate that TuMV induces root re-patterning. Plant viruses can disturb water management at a whole-organism scale, simultaneously modifying water loss rate through stomata and water intake from the roots (El Aou-ouad *et al.*, 2017). The importance of root-based events in plant-virus interactions has been pointed out already and future work should focus on assessing relevant root parameters under viral infections (Westwood *et al.*, 2013).

Additionally, viral factors interfering with stress-responsive genes can hamper adaptive drought responses, such as the down-regulation of RD29A detected here (Figure 6e, Figure 7e), or vice-versa: (Prasch and Sonnewald, 2013) found that the transcriptomic phenotype of TuMV-infected plants is different from the one that is triggered by drought or by the combination of both stresses, and proposed that specific combinations of abiotic or biotic stress act synergistically or antagonistically, respectively. These authors found also that transcripts related to defense genes were differently activated under the combination of TuMV + drought, and signaling components were reduced under multiparallel stress, probably deactivating TuMV-specific signaling networks, leading thus to increased susceptibility of plants under the combined stress.

Here, we concluded that TuMV reduced water loss through stomata in Arabidopsis but enhanced drought susceptibility. We found that stomatal closure correlated with enhanced total rosette ABA levels and highly increased transcription levels of ABA importer to guard cells ABCG40. Therefore, this work reinforces and expands previous findings concerning Arabidopsis-TuMV interaction regarding combined stresses (Prasch and Sonnewald, 2013) and hormone response (Manacorda *et al.*, 2013; Sánchez *et al.*, 2015; Casteel *et al.*, 2015; Poque *et al.*, 2018). Whereas recent reports studying combinations of virus plus drought stresses have pointed out to an increased tolerance to drought triggered by diverse viruses (Westwood *et al.*, 2013; Xu *et al.*, 2008; Aguilar *et al.*, 2017), the only previous report using Arabidopsis infected with TuMV (Prasch and Sonnewald, 2013) found increased susceptibility, consistent with the present work. All these reports, however, used the one-plant-per-pot approach, which has the shortcomings pointed by (Verslues *et al.*, 2006) regarding the assumption of equal root stress. Even (Xu *et al.*, 2008), who placed plants receiving different inoculation treatments (mock- or virus-inoculated) in the same trays, placed plants of each treatment group close together, thus creating a higher distance between roots of plants receiving different inoculation treatments. We chose to follow the approach of two plants per pot with “recovery after rewatering” proposed by (Verslues *et al.*, 2006) that allowed to quantify differences in survival rates between treatments with a simple χ^2^ test (Table 1). Whereas our work in any way invalidates those previous reports, we used an alternative methodological approach to study the combination of stresses that deal with challenges arising from differential stomatal aperture, root architecture and other traits affected by plant viruses. Apart from that, this work further reinforces the emerging concept that physiological outcomes and mechanisms in different plant-virus interactions are far from unique (Aguilar *et al.*, 2017).

## iv) Experimental procedures

### Plant growth conditions

*Arabidopsis thaliana* Col-0 and Ler ecotypes and mutant seeds were stratified at 4° C for 3 days and grown in a controlled environmental chamber (Conviron PGR14; Conviron, Winnipeg, Manitoba, Canada). For soil-based experiments, plants were grown in individual pots in trays and plants were well watered the day before measurements excluding drought experiments. For long days experiments, similar conditions as described by (Boyes *et al.*, 2001) were used, with minor modifications: A 15.5 hours light/7.5 hours dark photoperiod with T(°C) = 23/21, Hr(%) = 60/65, and a light intensity of 115 (+/-10) μE m^−2^ s^−1^ was used; additionally, two continuous ½ h ramps for T, Hr(%) and light intensity were added to simulate dawn/nightfall. For short days experiments (including all hydroponics experiments) growing conditions were identical except for photoperiod = 10/14. Except stated otherwise, all experiments were performed in short-days conditions. Hydroponic growth protocol, solutions and set-up were adapted from (Conn *et al.*, 2013).

### Virus infection assays

TuMV-UK1 strain (accession number X65978) (Sánchez *et al.*, 1998) and JPN1 strain (accession number KM094174) (Sánchez *et al.*, 2015) were maintained in infected *A. thaliana* Col-0. Fresh sap was obtained immediately prior to use in order to inoculate plants with sodium sulfite buffer (1% K_2_HPO_4_ + 0.1% Na_2_SO_3_ [wt/vol]). ORMV (Aguilar *et al.*, 1996) was maintained in *Nicotiana tabacum* (cv. Xhanti nn) and infective sap was obtained after grinding infected leaves with mortar and pestle in 50 mM phosphate buffer pH=7.5. Mock-inoculated plants were rubbed with carborundum dust with either sodium sulfite buffer or 50 mM phosphate buffer pH=7.5, respectively. Plants were mechanically inoculated at 1.08-1.09 growth stage (Boyes *et al.*, 2001) in true leaves #3 and #4 at around 22-24 days post-sowing. These leaves were almost fully developed by the time of the procedure and therefore constituted a source tissue for the export of molecular signals and virions systemically, particularly to those leaves forming closer angles with the inoculated ones (Farmer *et al.*, 2013; Manacorda *et al.*, 2013). To allow for inter-experiment and inter-parameter comparison, all measurements were performed at the same stage of viral infection (11-13 DPI) except otherwise stated. DPI = Days post-inoculation.

### Image acquisition

Infrared images were taken with an AVIO TVS-200P infrared camera (Nippon Avionics Co. Ltd., Japan) and analyzed with GORATEC software (©GORATEC Technology GmbH, Otto-Hahn-Strasse 21 D-85435, Erding, Germany). Infrared camera settings were adjusted according to the manufacturer’s User Manual. In order to facilitate the manual segmentation of the rosette from the otherwise more heterogeneous background in soil-based experiments, the substrate was covered with a porous black cloth. Similar T_leaf_ increases were registered relative to the mock-inoculated controls in both experimental setups. In experiments where specimens were imaged at different days post-inoculation (DPI), photographs were taken at the same time of the day on successive days. The average rosette temperature per plant (T_leaf_) was used for statistical analysis. Normalized ΔT_leaf_ between treatments was calculated setting average mock-inoculated rosette temperatures at 0 and then computing T_leaf_ differences for all individual plants. For overnight recording of infrared images, the infrared camera was placed standing on a tripod inside the growth chamber. Zenithal photographs of the tray containing hydroponically grown plants were taken at fixed intervals of time. Average rosette temperatures were obtained for each plant at each time point. A linear mixed model with treatment and time as fixed effects and plant as random effect was fitted to test for T_leaf_ temperature differences between treatments along all the time points (Table S1) and periods of day (Table S2). An autoregressive-moving average model of order 1 (ARMA(1, 1)) was found to be the best fit for both analyses. Visible-light images were taken with a Canon PowerShot SX50HS camera.

### Stomatal conductance and stomatal aperture analyses

Stomatal conductance (**g_s_**) measurements were performed with a SC-1 Leaf Porometer (METER Group, Inc. USA) following the manufacturer’s *Operator’s Manual Version 8* recommendations regarding calibration and general use. *G*_*s*_ was measured in the abaxial side of the middle part of leaf #8 across all treatments and genotypes at 24-27.9 °C and 46.7-66.9.% relative humidity (range of conditions across experiments) around noon. Stomatal aperture was measured in the abaxial side of the middle part of leaf #8 of infected or mock-treated plants. Two epidermal peels were used per leaf, and 20 stomata were randomly chosen for measurement in each peel. Peels were obtained using adhesive tape, and mounted for microscopic observation using a 10 mM KCl / 10 mM MES-KOH, pH = 6.15 solution. Four or five plants were analyzed per experiment and two independent experiments were performed, showing similar results. For both Col-0 and *abi2-1* data separate linear mixed effect models were applied, with treatment as fixed effect and individual (each one with n = 40 stomatal apertures measured) as random effect after checking that *Peel within Individual* had a negligible effect.

### Rosette water loss experiments

For relative daily water consumption, hydroponically grown plants were separately moved at 12 DPI from trays to closed Falcon tubes containing fresh BNS and weighted on a precision scale. Weight differences were recorded after 24 h. Rosette fresh weight (RFW) was measured to relativize water loss. Two independent experiments showing similar results were performed and the P-value (P = 0.0016) was calculated for the hierarchical ANOVA test of the linear model considering both experiments. For initial dehydration rate experiments, rosettes were detached at 12 DPI, imaged and placed on a precision scale (abaxial side upwards). Rosette weight loss was recorded every 15 s for 2 min. Taking into account the time-dependency of the data (serial correlation of data arising from measurements taken on the same experimental unit several times), differences in losses in relative rosette weight per projected area (PrA) per time regarding treatments were assessed using generalized least squares models. A comparison between several models led to the selection of a general correlation structure with different inter-time variances allowed. Multiple comparison tests were used to detect specific time points when differences between treatments were present.

### Hormone quantification

Individual plants growing in pots were harvested at 12 DPI and pooled within treatment (pool size 3 ≤ N ≤ 11 plants; five pools per treatment were used)). Plants were well-watered throughout the experiment and fully hydrated 20 h prior to harvest to avoid water status differences between treatments and to ensure no water-stress could affect hormone concentration determination. Systemic rosette tissue comprising leaves #8 and upper were harvested and immediately frozen in N_2_ (l). Extraction and purification of endogenous hormones was performed as described in (Durgbanshi *et al.*, 2005) and then, they were separated by reversed-phase HPLC, using an Alliance 2695 separation module (Waters; Milford, MA, USA) equipped with a Restek Roc C18 3 μm column (100 × 3.0 mm). The identification and quantification of hormones was performed using a quadrupole tandem mass spectrometer (Quattro Ultima, Micromass; Manchester, UK) fitted with an negative electrospray ion (ESI^−^) source, in multiple reactions monitoring mode (MRM). Source temperature was to 100°C. The collision energies and cone voltage were set to 15 eV and 35 V. The MassLynx spectrometry software program, V. 4.1 (Waters) was used for data analysis. Two independent experiments showing similar results were performed and p-values were calculated for the hierarchical ANOVA test of the linear model considering both experiments.

### Hormone treatment

SA at 0.5 mM (pH=7) was sprayed on Col-0 plants grown under LD conditions at growth stage 1.08. Control plants were sprayed with the same volume of the same solution, without SA (pH = 7). Effectiveness of treatment was confirmed by qRT-PCR using SA-inducible PR1 (AT2G14610) (Uknes *et al.*, 1992). Samples were harvested at 4 or 24 hours post treatment (HPT) and processed as described in (Manacorda *et al.*, 2013)

### .qPCR assays

Oligonucleotide primer sets for real-time (quantitative) PCR were designed using Vector NTI Advance 9 software (Life Technologies, Carlsbad, CA, U.S.A.). They are listed in Table S5. Details on the minimum information for publication of quantitative real-time PCR experiments (MIQE Guidelines: (Bustin *et al.*, 2009; Bustin *et al.*, 2010) are listed in Table S6. Primers were designed to target exon regions and for genes for which splice variants are predicted using ThaleMine, primers were designed in order to target all splice variants. For qPCR analysis, LinRegPCR program (Ramakers *et al.*, 2003) and the normalization method of (Pfaffl *et al.*, 2002) were used. Relative expression ratios and statistical analysis were calculated using fgStatistics software (Di Rienzo, 2009).

### Drought experiments

Seeds were sown either in a one or two plant-per-pot scheme (Verslues *et al.*, 2006). For each experiment, up to 24 pots were placed in a common tray. Treatments were randomly assigned to plants. In the one-plant-per-pot experiments, pots were mixed to minimize location effects. In the two plants per pot scheme one plant was randomly selected for TuMV infection and the other one was left as healthy control. Plants were fully bottom-watered to reach homogeneous pot weight just prior to inoculation (therefore 0 DPI = 0 DAWW). Afterwards, watering was withheld and plants were periodically imaged and inspected for development of symptoms of water stress. Then, plants were fully rewatered once from the bottom and the number of surviving plants was recorded after 3-5 days. The same experimental protocol was used for both long- and short-day photoperiods.

### Statistical analysis

Unless otherwise specified, all statistical analyses were performed using R (RStudio Team, 2016; RCoreTeam, 2016). For all statistical analysis, significance was set as: NS = P > 0.05, * = P ≤ 0.05, ** = P ≤ 0.01, and *** = P ≤ 0.001.

For data presenting nested (hierarchical) or serial correlation (time-dependency) structure, linear mixed effects models (*lme*) and generalized least squares models (*gls*) were implemented using the *nlme* package of R (Pinheiro *et al.*, 2020), used as suggested in (Pinheiro and Bates, 2000) for constructing, validating, comparing and selecting models. Multiple comparison tests were performed using the *emmeans* package (Lenth, 2019).

## (vii) Accession numbers

Arabidopsis mutants were in the Col-0 background except *abi2-1* (Ler). Those of them that were not a kind gift of fellow researchers were obtained from the Arabidopsis Biological Resource Center (http://www.arabidopsis.org). The accession numbers for the genes of the mutant plants used in this article are as follows: SID2 (At4g39030), NPR1 (At1g64280), AOS (At5g42650), LOX1 (At1g55020), ABA3 (At1g16540), ABI2 (At5g57050).

## (viii) Acknowledgements

We thank Dr. Flora Sánchez and Dr. Fernando Ponz for the kind gift of ORMV and TuMV virus (JPN1 and UK1 strains), Dr. Fernando Carrari for the lending of the Leaf Porometer (Decagon/METER), Dr. Rocío Tognacca for *abi2-1* seeds, Dr. Jean-Luc Montillet for *lox1-1* seeds, Dr. Annie Marion-Poll for *aba3-1* seeds, Lic. Pablo Cáceres for assistance in the use of the Genevestigator bioinformatics tool and technicians Agustín Montenegro, Matías Rodríguez and Ignacio Tévez for careful plant growth management.

## (ix) Short legends for Supporting Information

***Figure S1.*** TuMV induces an increase in T_leaf_ in *Arabidopsis thaliana* Col-0. ***(a,b)*** Images showing typical rosette phenotypes triggered by TuMV (UK1 strain) at 15 DPI under ***(a)*** visible and ***(b)*** infrared light (false color image with scale to the right). Average rosette temperature displayed. Both images display two representative plants for both mock-inoculated (upper row) and TuMV-infected (lower row) plants.

***Figure S2.*** TuMV induces similar phenotypes in hydroponics than in soil-based experiments and maintains reduced stomatal aperture during nighttime. ***(a)*** Hydroponics-grown plants display similar phenotypes than their soil-grown counterparts (15 DPI). ***(b)*** Infrared images were taken at several DPIs from 2-15 DPI, and separate t-tests were performed to assess T_leaf_ differences between treatments. N = 4 and n = 7 for mock and TuMV groups, respectively. ***(c-e)*** Hydroponics-grown plants were imaged from prior to dusk to after dawn, encompassing measurements for the entire nighttime between 12-13 DPI. ***(c)*** Representative IR photograph showing the tray with the plants. Green circles = mock-inoculated plants. Violet circles = TuMV-infected plants. Black box = plants not used for IR quantitative analysis. ***(d-e)*** Hydroponics-grown plants were imaged from prior to dusk to after dawn, encompassing measurements for the entire nighttime between 12-13 DPI. ***(d)*** Line graph showing group average rosette temperatures across the experiment for both treatments. ***(e)*** Box and jitter plot showing the data grouped by time of the day corresponding to the photoperiod set in the growing chamber. Note the dusk/dawn ramp.

***Table S1.*** ANOVA table showing the combined effect of treatment and time on T_leaf_ differences between hydroponics-grown plants.

***Table S2.*** ANOVA table showing the combined effect of treatment and time of day (= “period”) on T_leaf_ differences between hydroponics-grown plants.

***Table S3.*** Average T_leaf_ and ΔT_leaf_ between mock-inoculated and TuMV-infected plants during the different time of day (= “period”) categories. ΔT_leaf_ between treatments were found to be similar across different time of day (P > 0.05).

***Figure S3.*** TuMV induces a decrease in *g_s_* in hydroponic growth conditions. N = 8 per group.

***Figure S4.*** Effect of TuMV on stomatal conductance in mutants in genes related to hormone production and signaling.*G*_*s*_ measurements were taken in 12-DPI infected mutant plants and compared with healthy controls. All assays included measurements in Col-0 plants (infected vs. control), as WT controls, which always showed the low *g_s_* phenotype under TuMV infection (not shown).

***Figure S5.*** Reported expression pattern of *ABI2* in Arabidopsis. ***(a)*** Boxplots showing that *ABI2* is mostly expressed in guard cells, followed by root endodermis and senescent leaves. ***(b)*** Comparative expression of *ABI2* along 10 developmental stages (scatterplot). Note the overexpression in senescent tissues. ***(c-d)*** ABA is the most important perturbation inducing *ABI2* expression, with ABA-triggering stresses as drought, osmotic and salt stress following suite. ***(e)*** In the roots, NaCl strongly induces *ABI2* overexpression. For ***(c),*** a heat map shows changes in gene expression levels of *ABI2* across different experimental conditions as illustrated by Genevestigator V3. The log ratio (base 2) of the signal intensity relating treatment conditions to control conditions is illustrated using the color scale. For ***(d-e),*** “Electronic fluorescent pictographs” of *ABI2* are illustrated using the Arabidopsis eFP-Browser. Absolute signal intensities are illustrated using a color scale with low levels of expression colored yellow and high levels colored red.

***Table S4.*** Comparative analysis between reported gene expression summarized from Genevestigator V3 bioinformatic tools (first 4 rows) and qPCR results from this work (5^th^ to 7^th^ rows). Down-regulation of the genes analyzed in this work is shown in green for both main perturbations (stresses) reported by Genevestigator (first 2 rows) and qPCR results from this study (5^th^ to 7^th^ rows). Opposite pattern is shown in red. Non-significant differences found in this study are in gray. DR = Down-regulated; UR = Up-regulated; NS = Non-statistically significant changes in expression detected.

***Figure S6.*** Drought experiments with one plant per pot. ***(a), (b)*** Line graphs showing the kinetic of water loss (Ln scale) for pots with one plant only either ***(a)*** separately or ***(b)*** smoothed conditional means by treatment. ***(a)*** Interrupted lines indicate death plants before the ending of the experiment. ***(b)*** Confidence intervals (95%) are shadowed in gray. Starting from 18 DAWW, LnWeight differences between treatments were statistically significant (0.01<p<0.0001).

***Figure S7.*** SA application downregulates RD29A. QPCR analyses at 4 and 24 HPT (=hours post treatment). Bars = SE. N = 5 per group.

***Figure S8.*** ORMV, a phytovirus unrelated to TuMV, also increases rosette surface temperature. ***(a)*** Representative infrared image showing a mock-inoculated Arabidopsis Col-0 plant and three ORMV-infected plants exhibiting higher rosette surface temperature. ***(b)*** Box-plot graph showing T_leaf_ difference between treatments. N = 7 mock and 9 ORMV plants were imaged.

***Table S5.*** List of oligonucleotide primers used in qPCR experiments.

***Table S6.*** Minimum Information for Publication of Quantitative Real-Time PCR Experiments (MIQE Guidelines).

