## Supplementary material for "TuMV triggers stomatal closure but reduces drought tolerance in Arabidopsis": Table S1

ANOVA.ARMA.model

| | numDF | denDF | F-value | p-value |
|---|---|---|---|---|
| (Intercept) | 1 | 186 | 463591,53 | <.0001 |
| treatment | 1 | 6 | 35,29 | 0,00102 |
| t | 31 | 186 | 246,86 | <.0001 |
| treatment:t | 31 | 186 | 1,23 | 0,20104 |
