## Supplementary material for "TuMV triggers stomatal closure but reduces drought tolerance in Arabidopsis": Table S2

ANOVA.period.ARMA.model

| | numDF | denDF | F-value | p-value |
|---|---|---|---|---|
| (Intercept) | 1 | 242 | 568007,538 | <.0001 |
| treatment | 1 | 6 | 34,944 | 0,00104 |
| period | 3 | 242 | 488,364 | <.0001 |
| treatment:period | 3 | 242 | 0,480 | 0,69664 |
