## Supplementary material for "TuMV triggers stomatal closure but reduces drought tolerance in Arabidopsis": Table S3

Period.avr

| | treatment | period | Temp | ΔT |
|---|---|---|---|---|
| 1 | mock | day | 23,46 | |
| 2 | TuMV | day | 23,91 | 0,45 |
| 3 | mock | dusk | 22,37 | |
| 4 | TuMV | dusk | 22,72 | 0,36 |
| 5 | mock | night | 21,34 | |
| 6 | TuMV | night | 21,65 | 0,31 |
| 7 | mock | sunrise | 22,44 | |
| 8 | TuMV | sunrise | 22,82 | 0,38 |
