## Supplementary material for "TuMV triggers stomatal closure but reduces drought tolerance in Arabidopsis": Table S6

**Table S6.** Experimental conditions used for messenger RNA quantification by qRT-PCR assays based on MIQE requirements (Busti*n et a*l., 2009; Busti*n et a*l., 2010).

| **Experimental design** |  |
| --- | --- |
| Control groups | *A.thaliana* Col0 buffer-inoculated. |
| Treatment groups | *A.thaliana* Col0 inoculated with either JPN1/UK1 TuMV strains or ORMV. |
| **Sample** |  |
| Type of sample | Systemic rosette (leaves ≥ #8) or whole root tissue for hydroponically grown short day conditions plants / 11^th^ leaves (RD29A quantification in long day conditions in Figure S5). |
| Processing procedure | Liquid nitrogen homogenization |
| Sample frozen conditions | -80 ºC |
| Biological replicates | For systemic rosette (leaves ≥ #8) and whole root tissue (hydroponically grown short day conditions plants) 4 mock-inoculated and 7 UK1-infected plants were used. Rosette and root tissue were grinded separately. / Leaves #11 used for RD29A quantification were pooled when necessary to obtain enough tissue for RNA extractions (4dpi, 6≤n≤8; 10dpi, n=2-3 for each pool). Each pool was treated as a separate biological replicate and qRT-PCR experiments were performed using 4≤n≤7 pools for each treatment and DPI. |
| RNA: DNA-free | RT^-^ control without amplification. |
| **RNA extraction** |  |
| Procedure | Acid Phenol extraction |
| Reagents | TRIzol (Invitrogen) |
| Details of Dnase treatment | DNAse I Amp Grade (Invitrogen), 15 min at room temperature |
| Contamination assessment | < 3% |
| Nucleic acid quantification | Absorbance at 260 nm |
| Instrument and method | NanoDrop instrument |
| Purity( A260/ A 280) | > 2.01 (average, SD = 0.03) |
| Purity( A260/ A 230) | > 2.10 (average, SD = 0.28) |
| RNA integrity | Analyzed by agarose gel electrophoresis |
| **Reverse transcription** |  |
| Complete reaction conditions | Reaction was performed as described by the manufacturer´s instructions. |
| Amount of RNA and reaction volume | 1 µg of RNA, 20 µl |
| Priming oligonucleotide | Random primers (Invitrogen) |
| Reverse transcriptase | M-MLV (Invitrogen). |
| Temp and time | 10 min 50 ºC, 50 min 37 ºC, 15 min 70 ºC. |
| **qPCR protocol** |  |
| qPCR chemistry | SYBR green, ROX as passive reference. |
| Complete reaction conditions (*) | 5 min 95 ºC, (15 s 95 ºC, 30 s 60 ºC, 40 s 72 ºC) x 45 cycles |
| Reaction volume and amount of cDNA | 2 µl of a 1/20 dilution of synthesized cDNA in a final volume of reaction of 10 µl. |
| Primers, Mg^2+^ and dNTPs concentration | 3 mM Mg^2+^, 200 nM primers, 0,2 mM dNTPs |
| Polymerase | TransStart® TaqDNA Polymerase (TRANSGEN BIOTECH). |
| Buffer | 20 mM Tris-HCL (pH = 8.4), 50 mM KCl |
| Manufacturer of qPCR instrument | StepOnePlus, Thermo Fisher Scientific |
| **qPCR validation** |  |
| Specificity | Analysed by Melting Curve parameters on each qPCR run and automatic MTP assessment by StepOnePlus software (v2.3). NTC assessment. |
| Method of PCR efficiency calculation | Mean PCR efficiency per amplicon calculated by LingRegPCR program ((Ramaker*s et a*l., 2003)). |
| **Data analysis** |  |
| qPCR analysis program | LinRegPCR program |
| Method of Cq determination | LinRegPCR program |
| Outlier identification | LinRegPCR program |
| Justification of number and choice of reference genes | UBQ5 (NM_116090) was chosen as the most stable reference gene in our conditions based on our previous work (Manacord*a et a*l., 2013). |
| Description of normalization methods | (Pfaff*l et a*l., 2002)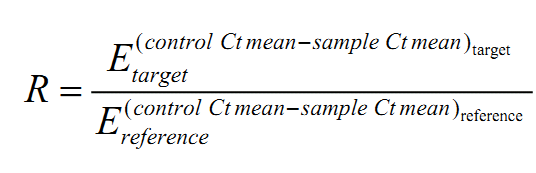 |
| Number of technical replicates | 2 |
| Statistical method | Permutation test |
| Software | fgStatistics software ((Di Rienzo, 2009) (<http://sites.google.com/site/fgStatistics/>) |
| Repeatability (intraassay variation) C_q_ mean SD error | Between 0.08 and 0.34 depending on the assayed amplicon. |

(*) Conditions are described for a generic qPCR assay. Particular annealing/extension temperatures could vary between amplicons. Specific conditions for each amplicon are available upon request.
